# Development of a multiparent advanced generation intercross (MAGIC) population for genetic exploitation of complex traits in *Brassica juncea*: glucosinolate content as an example

**DOI:** 10.1101/793331

**Authors:** Tianya Wang, Wei Wan, Kunjiang Yu, Aimal Nawaz Khattak, Botao Ye, Renqin Yang, Entang Tian

**Author notes:** These authors contributed equally to the work (Tianya Wang, Wei Wan).

## Abstract

Multiparent advanced generation intercross (MAGIC) populations have recently been developed to allow the high-resolution mapping of complex quantitative traits. This article describes the development of one MAGIC population and verifies its potential application for mapping quantitative trait loci (QTLs) in *B. juncea*. The population was developed from eight founders with diverse traits and composed of 408 F_6_ recombinant inbred lines (RILs). To develop one rapid and simplified way for using the MAGIC population, a subset of 133 RILs as the primary mapping population were genotyped using 346 intron-length polymorphism (ILP) polymorphic markers. The population lacks significant signatures of population structure that are suitable for the analysis of complex traits. Genome-wide association mapping (GWAS) identified three major glucosinolate (GSL) QTLs of *QGsl.ig01.1* on J01 for indole GSL (IG), *QGsl.atg09.1* on J09 and *QGsl.atg11.1* on J11 for aliphatic GSL (AG) and total GSL (TG). The candidate genes for *QGsl.ig01.1, QGsl.atg09.1* and *QGsl.atg11.1* are *GSH1, GSL-ALK* and *MYB28*, which are involved in converting glutamate and cysteine to *γ*–EC, the accumulation of glucoraphanin, and the whole process of AG metabolism, respectively. One effective method for association mapping of quantitative traits in the *B. juncea* MAGIC population is also suggested by utilization of the remaining 275 RILs and incorporation of the novel kompetitive allele specific PCR (KASP) technique. In addition to its QTL mapping purpose, the MAGIC population could also be potentially utilized in variety development by breeders.

## Introduction

With the rapid development of next-generation sequencing technologies and the dramatic decrease in the cost of sequencing, the detection of quantitative trait loci (QTLs) in plant breeding is no longer limited by the availability of genetic marker information and genotyping throughput but rather by the genetic materials employed [1]. Traditional QTL mapping populations combine the genomes of two parents and can only capture a small snapshot of the genomic regions differing between the two parents. While the alternative of associating mapping has far greater diversity and sufficient power to detect genomic regions of interest, it needs a very large population and may have difficulty detecting rare alleles of importance [2].

In recent years, we have witnessed the rise of multiparent populations with a new type of experimental design, offering great advantages for genetic studies in plants. The multiparent advanced generation intercross (MAGIC) population is one of the most popular multiparent populations, in which the multiple inbred founders are intercrossed several times to combine all the genomes of the founders in a single line, resulting in a diverse population whose genomes are fine-scale mosaics of the contributions from all founders [3]. It generates a diverse population whose genomes recombine many times, which is suitable for high-resolution trait mapping [2]. In addition, MAGIC populations can also provide excellent materials for plant breeding due to their features of high recombination and the resulting diverse phenotypic diversity. The term MAGIC was first coined in 2007 [4], and the first MAGIC population in plant species was developed in *Arabidopsis thaliana* (L.) [5]. Then, dozens of MAGIC populations were established in a wide range of various crops, including rice [6–10], tomato [11, 12], fava bean [13], cowpea [14], maize [15], barley [16, 17], strawberry [18], sorghum [19], and wheat [3, 20–23]. However, no MAGIC populations in Brassicaceae have been reported to our knowledge.

To verify the suitability of the newly developed *B. juncea* MAGIC population for gene mapping, several glucosinolate (GSL) traits were chosen for further association analysis using the population. GSLs were first discovered in mustard seeds during an exploration of the chemical origin of their sharp taste in the 17^th^ century. In the Brassicaceae family, GSLs are the major secondary metabolites and can be synthesized in all species of this family via a three-part biosynthetic pathway from methionine, tryptophan and phenylalanine [24–26]. GSLs are mainly divided into aliphatic, indolic and benzyl GSLs in *Brassica* species. As important secondary metabolites, GSLs can be hydrolyzed by myrosinase to form different toxins, such as nitriles, isothiocyanates, and thiocyanate. These toxins have been widely studied in regard to their function in altering insect resistance [27, 28] and herbivory [29–31], nematode survival [32, 33], and microbial pathogens [34, 35]. More importantly, certain GSLs might have potential anticarcinogenic activity in mammals [36–38]. The importance of GSLs makes them a very special complex trait, and their metabolic mechanism is well studied in *A. thaliana*. More work is needed in *Brassica* crops in regard to their polyploidy feature.

*Brassica juncea* exhibited better drought and heat tolerance, disease resistance, insect resistance and shattering resistance than *B. napus* [39–45]. In addition to its use as a condiment in Canada and China and as a vegetable in China, great efforts have been made to develop *B. juncea* as an alternative oilseed crop, especially in semiarid areas. In the present study, we try to develop and use one MAGIC population to exploit the complex traits in *B. juncea*. For genotype detection, we used ILP markers, which were designed in exons flanking the target intron, which is highly polymorphic, specific, codominant, neutral, convenient and reliable [46, 47]. ILP-type markers have been used for genotyping studies in many species, such as in rice, yellow mustard, foxtail millet, maize, tomato, *B. juncea*, *B. rapa* and *A. thaliana* [46–52]. The genotyping results are used to estimate the population structure, in linkage disequilibrium (LD) analysis, and in genome-wide association study (GWAS) analysis. The detected QTLs for GSLs demonstrate that the MAGIC population is an excellent platform for exploiting the complex traits in *B. juncea*.

## Materials and Methods

### Plant materials and field experiment

The eight founders of the MAGIC population were chosen from 34 *B. juncea* germplasm lines in our earlier field experiments [53]. Two rows were planted for each line of the 408 lines and the eight founders in the MAGIC population in 2018 and 2019, respectively. All the materials were planted in the field of Guizhou University, Guiyang, China. Three self-pollinated plants were randomly chosen to harvest seeds for GSL content analysis in 2018 and 2019, respectively.

### GSL component analysis

The 2-propenyl and 3-butenyl GSL contents of the mature seeds from each plot were analyzed following published methods [54, 55] with minor modifications. Each seed sample was crushed, and 200 mg of each sample was extracted twice with 2 ml boiling 70% methanol. The concentration of GSLs in the seeds was determined by high-performance liquid chromatography (Waters 2487/600/717) using the ISO9167-1 (1992) standard method. The AG is the sum of 2-propenyl GSL, 3-butenyl GSL and 4-pentenyl GSL. The TG is the sum of AG, IG and 4-hydroxy-3-indolylmethy GSL (HOI GSL).

### ILP analysis

Genomic DNA was extracted from young leaves of eight founders and the 408 MAGIC lines using the modified sodium dodecyl sulfate method [56]. PCR of the ILP markers was carried out according to our previous studies [57, 58]. Each PCR (20 μl) contained 1× standard PCR buffer (NEB), 1 U of Taq polymerase (NEB), 0.25 μM forward primer, 0.25 μM reverse primer, 100 μM each dNTP and 50 ng of genomic DNA in a total volume of 20 μL. The PCR amplification consisted of an initial denaturation step at 94°C for 5 min; 35 cycles consisting of 94°C (45 sec), 55°C (45 sec) and 72°C (1 min); followed by termination at 72°C for 7 min. All PCR products were analyzed by electrophoresis in 2% agarose gels in 1× tri-acetate-ethylene diamine tetra acetic acid buffer. Gels were visualized by staining in ethidium bromide and photographed on a digital gel documentation system.

### Population structure and LD analysis

PCA was used to assess the MAGIC population using the genotyping data, which was performed in SPSS 20.0. The first two principal components (PCs) were used to visualize the dispersion of the lines within a population in a graph. Pairwise similarity coefficients were calculated for all pairwise combinations within a population according to the method developed by (Nei and Li) implemented in the software TASSEL v5.0 [59]. The neighbor-joining (NJ) tree based on this genetic similarity matrix was constructed accordingly. To display the clustering feature, clustering analysis was conducted in R. The linkage disequilibrium (LD) analysis of the MAGIC population was performed by computing the *r*^*2*^ values between pairs of ILP markers using Tassel v3.0 [55]. A significance threshold of p< 0.001 was used to screen the LD locus pairs, and the remaining pairs were not considered informative. The parameter *r*^*2*^ was used to graphically represent the LD curves with the R software.

### Marker-trait association analyses

Marker-trait association analyses between the three GSL quality traits of two consecutive years and the ILP genotyping data were conducted using a general linear model (GLM) with the TASSEL 3.0 software package [59]. The significance threshold for the associations between ILP markers and traits was set as p< 0.001 (−log10(p) = 3). The ILP markers prefixed with “At” were mapped on the chromosomes of the *B. juncea* genome according to Panjabi’s genetic map [52]. In addition, the ILP markers prefixed with “Bnap” and “Brap” were mapped on the chromosomes of the *B. juncea* genome by blasting their originated DNA sequence with the *B. juncea* genome (Brassica_juncea_v1.5,http://brassicadb.org/brad/datasets/pub/Genomes/Brassica_juncea/V1.5/Bju.genome.fasta.gz) and assigned a genome location using a BLAST-like alignment tool [60].

### Data availability

The MAGIC lines and founders are maintained by Institute of Oil Crops of Guizhou University (IOC-GZU) and are available upon request. S1 Figure contains supplemental figures and tables cited in this article. S1 Table contains the information of the distribution of ILP type markers in different chromosomes. S2 Table contains the data of GSL contents (umol/g) of the PAM-MAGIC population in 2018 and 2019. Supplemental File 1, File 2 and File 3 contain genotype, phenotype and population sturcture for each MAGIC line used for GWAS analysis, respectively.

## Results

### Development of the MAGIC population

The initial *B. juncea* MAGIC population was developed by intercrossing 8 founder lines that are diverse in agronomic and quality traits (Table 1). The first stage followed a half-diallel mating system by intermating the 34 Chinese native *B. juncea* germplasm lines, and 17 biparental crosses were performed. The agronomic and quality traits of the 34 germplasm lines were estimated in our earlier study [53], and the eight diverse lines and their derived 4 F_1_s were selected for the next step. The resulting 4 F_1_s were intercrossed to derive 4-way crosses for which 2 such 4-way crosses integrated the four founders’ genomes of ABCD and EFGH. The last stage involved intercrossing the 2 4-way crosses to derive one 8-way cross integrating the eight founders’ genomes of ABCDEFGH. Finally, 600 seeds of the 8-way crosses were self-pollinated to produce the final MAGIC population including 408 F_6_ RILs with single-seed descent methods. The crossing scheme for the *B. juncea* MAGIC populations is shown in Fig 1.

**Table 1.** Features of the eight founder lines in developing the *B. juncea* MAGIC population.

| Code | Name | Features |
| --- | --- | --- |
| P<sub>A</sub> | 7H881 | larger seed size, low number of grains per pod, low total GSL content |
| P<sub>B</sub> | YufengZC | larger seed size, shorter, high total GSL content |
| P<sub>C</sub> | K100 | taller, early flowering, small seed size, |
| P<sub>D</sub> | SL105 | taller, early flowering, small seed size, long length of main inflorescence, thinner stem perimeter |
| P<sub>E</sub> | SL14 | thinner stem perimeter, taller |
| P<sub>F</sub> | ShilunYC | taller, more number of primary branches, bigger number of grains per pod |
| P<sub>G</sub> | LengjiaoYC | shorter, small seed size |
| P<sub>H</sub> | T6342 | shorter, smaller number of grains per pod |

**Fig 1.**
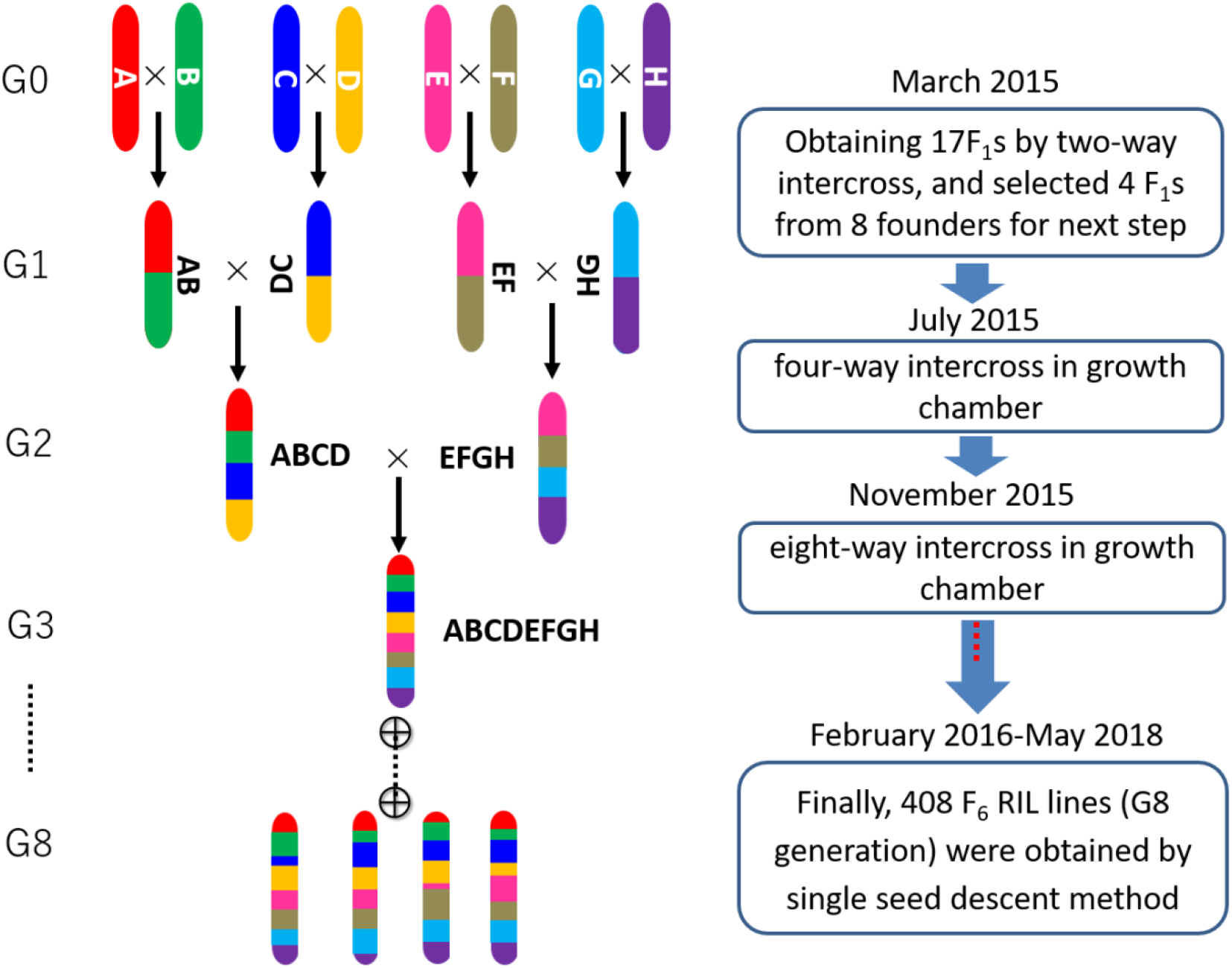
The scheme and flow of the MAGIC population development. Eight founders: A: 7H881, B: YufengZC, C: K100, D: SL105, E: SL14, F: ShilunYC, G: LengjiaoYC, H: T6342.

### Amplification fragment polymorphisms of the ILP markers

To increase the working efficiency, the MAGIC population is divided into two subpopulations for primary association mapping (PAM) and subsequent fine association mapping (FAM). The PAM-MAGIC and the FAM-MAGIC populations include 133 and 275 randomly selected lines, respectively. In this study, the genetic structure of the PAM-MAGIC population was analyzed to estimate the genetic structure of the whole MAGIC population. A total of 1,272 ILP primers, 284 from *Arabidopsis thaliana* [52], 745 from *B. napus* and 243 from *B. rapa* available in the Potential Intron Polymorphism (PIP) database [46], were used to screen the eight founders for polymorphic primers.

Of the 1,272 ILP primers, 296 (23.3%) generated clear and scorable polymorphic bands, varying in size from 150 to 1250 bp, among the eight founders. The 296 primers were used for genotyping the PAM-MAGIC population and produced a total of 346 polymorphic markers for estimation of the subpopulation’s genetic structure. The polymorphic markers are distributed evenly across the whole *B. juncea* genome (S1 Table), as reported in our previous publications [57, 58, 61].

### Population structure and LD analysis

Neighbor-joining tree and PCA methods are introduced to estimate the genetic structure of the PAM-MAGIC population. Fig 2A and 2B show a dendrogram from this cluster analysis. The eight founder cultivars are distributed almost evenly in the dendrogram, although 7H881 and SL105 were closer to each other than to the other cultivars. Fig 2C shows the proportion and the cumulative proportion of all the PCs derived from the PCA. The proportions of variance of PC1 and PC2 are 6.26% and 5.33%, respectively. The scatter plot made with PC1 and PC2 indicates that the distribution of the PAM-MAGIC lines is almost even and not clustered (Fig 2D).

**Fig 2.**
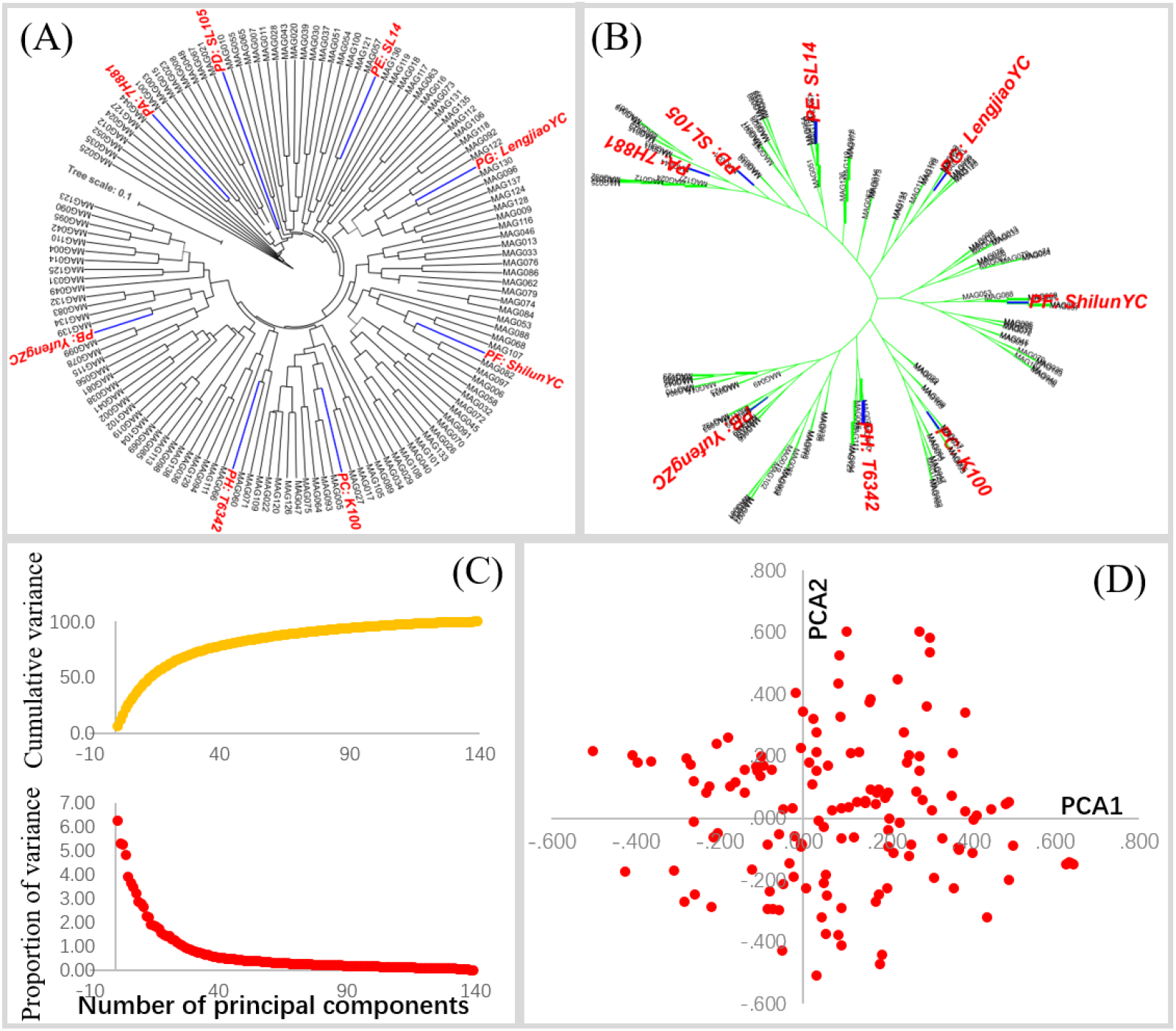
Population structure analysis through clustering (A and B), PCA (C) and a scatter plot (D).

LD analysis was conducted to explore the genome landscape of the PAM-MAGIC population for association studies. The squared Pearson correlation coefficients (*r*^*2*^) and D’ (measures of LD) were measured according to these 346 ILP polymorphic markers to describe the LD relationship between markers. The LD indicates a very low level in PAM-MAGIC lines (S1 Figure). A total of 59,686 possible pair combinations were obtained with an average of *r*^*2*^ = 0.034. The distribution of *r*^*2*^ values was concentrated in the < 0.1 interval that accounted for 95.3% of all pairwise coefficients, and 58.2% of the coefficients were < 0.01. The distribution of D’ values differed, with approximately 5% and 42.8% of the values being between 0.1 and 0.3. In addition, only 4.46, 5.16, and 8.11% of the coefficients (*r*^*2*^) were revealed to be significant at p < 0.0001, p < 0.001 and p < 0.01, respectively.

### Performance of phenotype and QTL mapping

To verify the suitability of our new *B. juncea* MAGIC population for gene mapping, as predicted from our genetic analysis, we conducted GWAS for the GSL traits of IG, AG and TG. Phenotypic dissection and depiction of genetic variability from a 2-year GSL dataset is summarized in S2 Table. The GSL contents were stable and significantly correlated for IG (0.987), AG (0.917) and TG (0.920) between 2018 and 2019. The broad-sense heritability of IG, AG and TG are 82.6%, 96.9% and 97.1%, respectively.

The associations between ILP markers and the GSL traits were analyzed with a naive general linear model (GLM) in Tassel 3.0 [59]. The markers from the same chromosomes with lower significance thresholds of p < 0.001 (−log10(p)=3) are used to analyze the real QTL region. To determine the confidence interval for the real QTL identified in this study, markers significantly associated with the GSL content that were colocated together and/or adjacent within intervals inferior to the LD decay distance of up and down region from the association peak were considered as being in the same QTL region. Finally, three QTL regions for IG, AG and TG in 2018 and 2019 were identified (Fig 3). Each QTL in one year is mapped onto Panjabi’s genetic map [52] and the physical map of *B. juncea* (Fig 3). The *B. juncea* physical map was constructed by blasting the DNA sequence of ILP markers prefixed with BnapPIP and BrapPIP to the *B. juncea* genome. The overlapped QTL regions of At4g37510 (R^2^=20.2% and 19.6%) and BnapPIP1698 (R^2^=21.1% and 20.5%), for IG, are mapped onto J01 and renamed *QGsl.ig01.1* (Table 2). The overlapped QTL regions of At5g66290 (R^2^=23.3 and 22.0 for AG and R^2^=23.4 and 21.1 for TG) and BnapPIP1437 (R^2^=10.1 and 7.9 for AG and R^2^=10.3 and 9.5 for TG), for AG and TG, are mapped onto J11 and renamed *QGsl.atg11.1* (Table 2). In addition, another overlapped QTL regions of At5g63905 (R^2^=22.9 and 18.9 for AG and R^2^=23.0 and 19.9 for TG) and BnapPIP1490 (R^2^=26.2 and 25.0 for AG and R^2^=26.4 and 24.1 for TG), for AG and TG, are mapped onto J09 and renamed *QGsl.atg09.1* (Table 2).

**Table 2.** The QTLs obtained from association mapping of IG, AG and TG.

| GSLs | QTL Name | Markers | R<sup>2</sup> (%)<sup>1</sup> | -LOG<sub>10</sub>P | Chr<sup>2</sup> | Position<sup>3</sup> | Consensus QTL name | Candidate Genes |
| --- | --- | --- | --- | --- | --- | --- | --- | --- |
| IG | IG01-18 | At4g37510 | 20.2 | 7.5 | J01 | 6.4cM | *QGsl.ig01.1* | *GSH1* [63-66] |
|  | IG01-19 | At4g37510 | 19.6 | 7.3 | J01 | 6.4cM |  |  |
|  | IG01-18' | BnapPIP1698 | 21.1 | 7.9 | J01 | 4,290,074bp |  |  |
|  | IG01-19' | BnapPIP1698 | 20.5 | 7.7 | J01 | 4,290,074bp |  |  |
| AG | AG11-18 | At5g66290 | 23.3 | 8.5 | J11 | 51.2cM | *QGsl.atg11.1* | *Myb28* [67-69] |
|  | AG11-19 | At5g66290 | 22.0 | 8.1 | J11 | 51.2cM |  |  |
|  | AG11-18' | BnapPIP1437 | 10.1 | 3.7 | J11 | 1,500,610bp |  |  |
|  | AG11-19' | BnapPIP1437 | 7.9 | 3.0 | J11 | 1,500,610bp |  |  |
|  | AG09-18 | At5g63905 | 22.9 | 8.1 | J09 | 30.1cM | *QGsl.atg09.1* | *GSL-ALK* [70, 71] |
|  | AG09-19 | At5g63905 | 18.9 | 6.8 | J09 | 30.1cM |  |  |
|  | AG09-18' | BnapPIP1490 | 26.2 | 9.8 | J09 | 10,280,466bp |  |  |
|  | AG09-19' | BnapPIP1490 | 25.0 | 9.4 | J09 | 10,280,466bp |  |  |
| TG | TG11-18 | At5g66290 | 23.4 | 8.5 | J11 | 51.2cM | *QGsl.atg11.1* | *Myb28* [67-69] |
|  | TG11-19 | At5g66290 | 21.1 | 7.8 | J11 | 51.2cM |  |  |
|  | TG11-18' | BnapPIP1437 | 10.3 | 3.7 | J11 | 1,500,610bp |  |  |
|  | TG11-19' | BnapPIP1437 | 9.5 | 3.5 | J11 | 1,500,610bp |  |  |
|  | TG09-18 | At5g63905 | 23.0 | 8.2 | J09 | 51.2cM | *QGsl.atg09.1* | *GSL-ALK* [70, 71] |
|  | TG09-19 | At5g63905 | 19.9 | 7.2 | J09 | 51.2cM |  |  |
|  | TG09-18' | BnapPIP1490 | 26.4 | 9.9 | J09 | 10,280,466bp |  |  |
|  | TG09-19' | BnapPIP1490 | 24.1 | 9.1 | J09 | 10,280,466bp |  |  |

<sup>1</sup>: Percentage of the total phenotypic variation explained by the QTL.
<sup>2</sup>: Abbreviation of chromosomes.
<sup>3</sup>: The position from the genetic map with a postfix of cM [52] and the physical position with a postfix of bp obtained by blasting the DNA sequence of ILP markers against the *B. juncea* genome [60].

**Fig 3.**
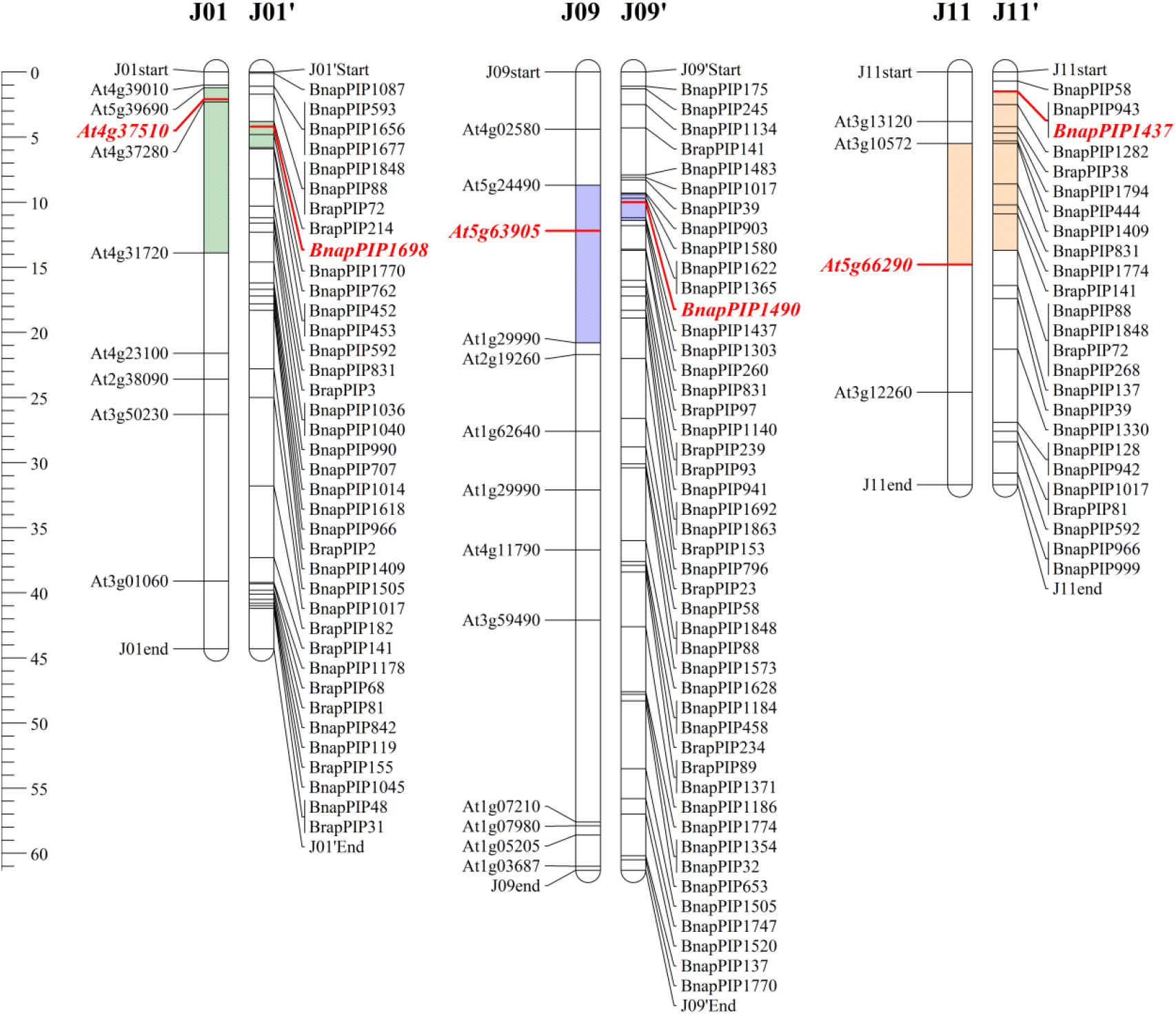
QTLs detected on *B. juncea* chromosomes J01, J09 and J11 for IG, AG and TG. The physical maps of J01’, J09’ and j11’ were constructed by blasting the sequence of PIP markers against the *B. juncea* genome. The chromosomes of J01, J09 and J11 are based on the published genetic map [1], and the physical position of each marker was deduced according to the published *B. juncea* genome [2]. The physical positions of the consensus QTLs were makered on different chromosomes using different colors. The unit of the ruler on the left is Mb.

### Candidate genes that may be associated with GSL synthesis

The physical regions of consensus *QGsl.ig01.1* on J01, *QGsl.atg11.1* on J11 and *QGsl.atg09.1* on J09 are located at 3,924,530-5,949,433 bp, 5,668,412-14,027,300 bp and 9,541,599-1,164,980 bp, respectively (Fig 3). The sequences of these two QTL regions were downloaded and utilized to predicate candidate genes using the annotated *B. juncea* reference sequence (*Brassica_juncea_v1.5*) [62]. The candidate gene of *QGsl.ig01.1* is *GSH1*, which is responsible for converting glutamate and cysteine to *γ*–EC [63–66]. For the consensus QTL of *QGsl.atg11.1*, the candidate gene is *Myb28*, which is involved in the whole process of aliphatic GLS (AG) metabolism *Myb28* [67–69]. Moreover, the candidate gene of *QGsl.atg09.1* is *GSL-ALK*, which is responsible for the accumulation of glucoraphanin [70, 71].

## Discussion

Dozens of studies have reported the utilization of MAGIC populations in many crops, such as rice and wheat. However, no such population has been reported for Brassicaceae. This paper provides another alternative for QTL mapping in addition to traditional biparental mapping populations and diverse germplasm populations in Brassicaceae. Within comparison to the MAGIC population, which has a higher number of parents and recombination events, the biparental population has limited genetic diversity and complexity. Even though the genetic complexity and diversity of the natural population is higher than that of the MAGIC population, its confounding population structure [72–74] enhances the risk of detecting false positives [75–77]. While the population structure of MAGIC is ignorable, the population structure obtained in this study is suitable for QTL mapping. Thus, to date, the MAGIC mapping population is the most ideal mapping population.

To verify the potential application of the MAGIC mapping population in mapping QTLs in *B. juncea*. We selected the three GSL traits of IG, AG and TG as candidates for association mapping. In this study, we mapped the three QTLs of *QGsl.ig01.1* for IG, *QGsl.atg11.1* and *QGsl.atg09.1* for AG and TG through association mapping. The three candidate genes are well studied in *Arabidopsis thaliana* and less studied in other plants. *GSH1* is reported to be responsible for converting glutamate and cysteine to *γ*–EC in *A. thaliana* [63–66]. *GSH1* is not specific to IG biosynthesis, which is involved in the biosynthesis pathway of all GSLs. Therefore, some other types of regulation, such as methylation, RNA and protein levels, might function and result in the phenotypic variation of IG content in the MAGIC population. As an important transcription factor, *Myb28* is specific for regulating the whole process of aliphatic GLS (AG) metabolism [67–69]. Moreover, *GSL-ALK* corresponding to the QTL of *QGsl.atg09.1* is responsible for the accumulation of glucoraphanin [70, 71]. Specifically, *GSL-ALK* can convert 4-methyl sulfinyl butyl and 4-methyl sulfinyl pentyl to 3-butenyl GSLs and 4-pentenyl GSLs, respectively [65, 78, 79]. The 3-butenyl GSLs and 4-pentenyl GSLs are the main individual components of AG, and AG is the main component of TG. Therefore, *GSL-ALK* and *MYB28*, corresponding to QTLs of *QGsl.atg09.1* and *QGsl.atg11.1*, respectively, must be the key factors involved in the accumulation and phenotyptypic variation of AG and TG in *B. juncea* MAGIC population in the present study. Moreover, the IG content is also one of the individual components of AG and TG, while the QTL of *QGsl.ig01.1* for IG was not detected when performing association mapping analysis for AG and TG. The main reason should be the small ratio of the IG content in AG and TG. This work can provide useful information for the mechanism of GSL biosynthesis and molecular marker-assisted breeding (MAS); however, more work is needed to clone and verify the function of these two genes in the future.

The *B. juncea* MAGIC population provided a template for the development and utilization of this type of population in other species of Brassicaceae. MAGIC populations have been constructed because of their attractive combination of gene discovery and breeding utility for geneticists and breeders. For breeders, the MAGIC population showed wider phenotypic variation in almost all traits than the eight founders [8], as observed in this study, which suggests the formation of transgressive segregants in the breeding direction. Interestingly, one breeding program in IRRI was started, which includes 700 lines from a total of 21 high-yielding lines [8]. Moreover, high-throughput SNP genotyping platforms are becoming cheap, and statistical approaches that can be applied to such populations are now available [8]. In this study, we used one subset of 133 randomly chosen RILs to estimate the population structure and conduct primary association mapping. The population was used to successfully map two QTLs for IND, AG and TG contents. The FAM-MAGIC population with the remaining 275 F_6_ lines is not useless and could be used for fine mapping through the utilization of the kompetitive allele specific PCR (KASP) technique for more complex quantitative traits. The KASP technique has been developed and successfully used for the fine mapping of the *Cf-10* gene in tomato through the utilization of bulk sequencing and the subsequent use of the KASP technique [80]. The KASP technique is useful in QTL mapping, especially for joint analysis by fully utilizing the genetic recombinant information of multiple populations. For the association mapping of more complex agronomic traits, the marker density should be increased by the utilization of the resequencing technique in case some minor QTL information is missed in the future. Then, the MAGIC population could play a greater role in gene discovery and breeding in *B. juncea*.

## Conclusion

In this study, we reported the first MAGIC population developed in Brassicaceae. The population structure and LD analysis indicate that the population is suitable for association mapping. To verify the usefulness of the MAGIC population, the association mapping of IG, AG and TG were analyzed. Finally, one QTL of *QGsl.ig01.1* for IND and two overlapping QTLs of *QGsl.atg09.1* and *QGsl.atg11.1* for AG and TG were identified, all of which play an important role in GSL metabolism. Thus, the MAGIC population provides an excellent planform for gene discovery and breeding in *B. juncea*.

## Acknowledgments

This work was funded by National Natural Science Foundation of China (Grant No. 31560422), Agricultural Science and Technology Support Program of Guizhou Province (Qiankehe zhicheng No. [2019]2396), Science and Technology Foundation of Guizhou Province of China (Grant No. Qiankehe J zi [2015]2052), Scientific Research Foundation for Returned Scholars, Ministry of Education of China (Grant No. Jiaowaisiliu [2015]1098), Construction Program of Biology First-class Discipline in Guizhou (Grant No. GNYL[2017]009).

## Supplementary materials

**S1 Table.** The distribution of ILP type markers in different chromosomes.

| Chromosomes | ILP markers from *B. napus* and *B. rapa*<sup>1</sup> | ILP markers from *A. thaliana*<sup>2</sup> | Total |
| --- | --- | --- | --- |
| J01 | 39 | 8 | 47 |
| J02 | 16 | 10 | 26 |
| J03 | 44 | 14 | 58 |
| J04 | 28 | 3 | 31 |
| J05 | 36 | 6 | 42 |
| J06 | 38 | 6 | 44 |
| J07 | 22 | 4 | 26 |
| J08 | 42 | 10 | 52 |
| J09 | 46 | 14 | 60 |
| J10 | 31 | 4 | 35 |
| J11 | 29 | 5 | 34 |
| J12 | 33 | 16 | 49 |
| J13 | 36 | 5 | 41 |
| J14 | 30 | 6 | 36 |
| J15 | 41 | 2 | 43 |
| J16 | 27 | 4 | 31 |
| J17 | 35 | 5 | 40 |
| J18 | 38 | 5 | 43 |
| Total | 611 | 127 | 738<sup>3</sup> |

<sup>1</sup>: The marker position of these markers prefixed “Bnap and Brap” is mapped onto *B. juncea* genome by the BLAST-like alignment tool [60].
<sup>2</sup>: The marker position of these markers prefixed “At” is from the published *B. juncea* genetic map [52].
<sup>3</sup>: Some primers are mapped onto more than one chromosome.

**S2 Table.** Glucosinolate contents (umol/g) of the PAM-MAGIC population in 2018 and 2019.

| Traits | Year | Mean | SD<sup>1</sup> | Range | CV (%)<sup>2</sup> |
| --- | --- | --- | --- | --- | --- |
| IG | 2018 | 0.62 | 0.51 | 0.10~2.62 | 82.26 |
| IG | 2019 | 0.56 | 0.51 | 0.00~2.62 | 91.07 |
| AG | 2018 | 101.33 | 33.54 | 28.98~197.91 | 33.10 |
| AG | 2019 | 103.32 | 38.67 | 28.48~218.76 | 37.43 |
| TG | 2018 | 106.75 | 33.71 | 33.91~204.44 | 31.58 |
| TG | 2019 | 106.79 | 38.22 | 31.07~211.82 | 35.79 |

<sup>1</sup>: SD: standard deviation.
<sup>2</sup>: CV: coefficient of variation, CV=SD/Mean

**S1 Figure.**
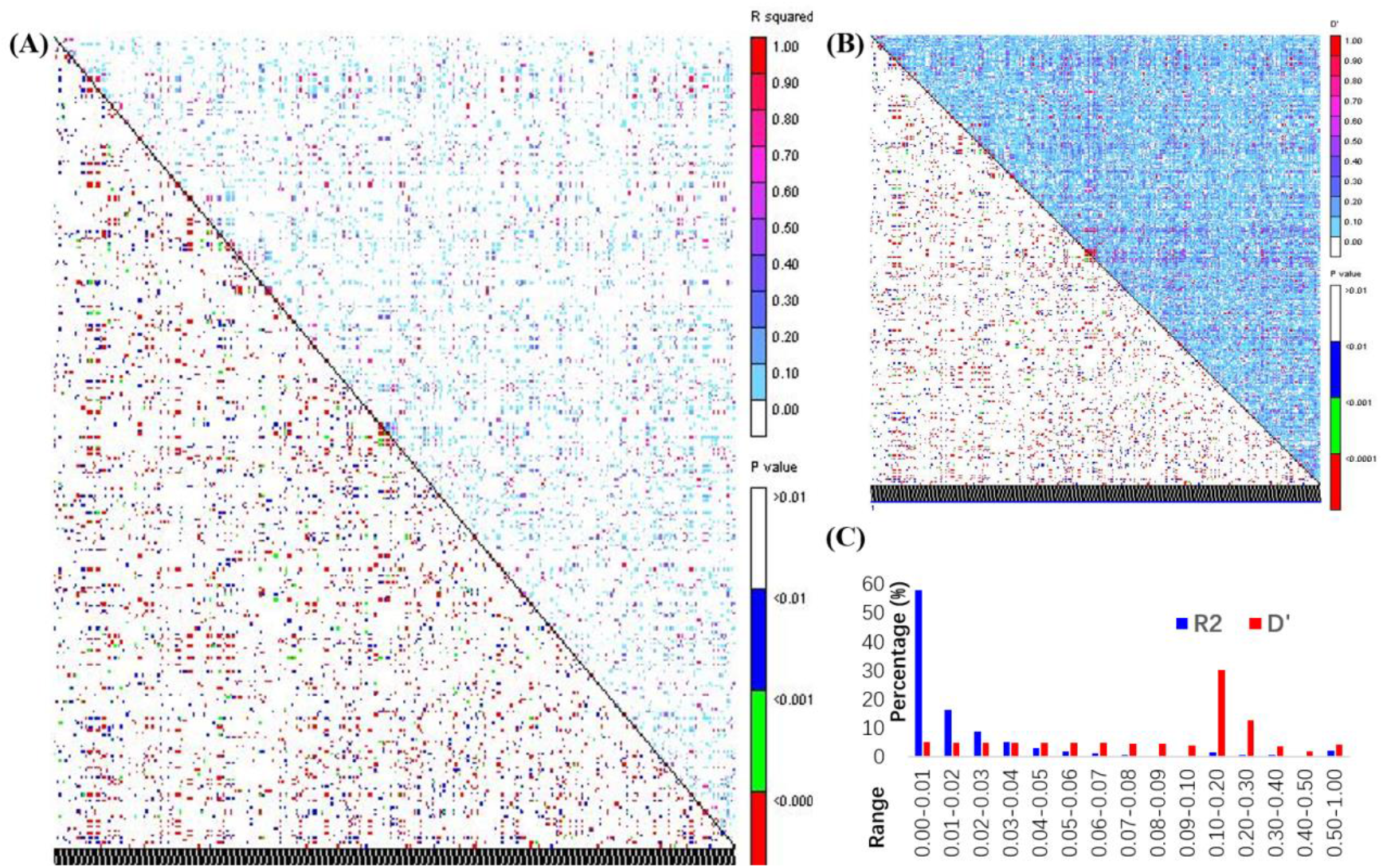
Linkage disequilibrium (LD) analysis for exploring the genome landscape of the PAM-MAGIC population for association studies. The squared Pearson correlation coefficients (*r*^*2*^, A) and D’ (measures of LD, B) are measured and analyzed (C).

